# Efficient Development of Platform Cell Lines Using CRISPR-Cas9 and Transcriptomics Analysis

**DOI:** 10.1101/2020.09.16.299248

**Authors:** Andrew Tae-Jun Kwon, Kohta Mohri, Satoshi Takizawa, Takahiro Arakawa, Maiko Takahashi, Bogumil Kaczkowski, Masaaki Furuno, Harukazu Suzuki, Shunsuke Tagami, Hidefumi Mukai, Erik Arner

## Abstract

Antibody-drug conjugates offers many advantages as a drug delivery platform that allows for highly specific targeting of cell types and genes. Ideally, testing the efficacy of these systems requires two cell types to be different only in the gene targeted by the drug, with the rest of the cellular machinery unchanged, in order to minimize other potential differences from obscuring the effects of the drug. In this study, we created multiple variants of U87MG cells with targeted mutation in the *TP53* gene using the CRISPR-Cas9 system, and determined that their major transcriptional differences stem from the loss of p53 function. Using the transcriptome data, we predicted which mutant clones would have less divergent phenotypes from the wild type and thereby serve as the best candidates to be used as drug delivery testing platforms. Further *in vitro* and *in vivo* assays of cell morphology, proliferation rate and target antigen-mediated uptake supported our predictions. Based on the combined analysis results, we successfully selected the best qualifying mutant clone. This study serves as proof-of-principle of the approach and paves the way for extending to additional cell types and target genes.

## 1 Introduction

Targeted drug delivery through the use of monoclonal antibodies has been a prominent therapeutic research area due to its promise of avoiding negative drug side effects by targeting diseased cells expressing specific antigens (Scott et al., 2012). This can be especially beneficial in cancer treatment, as the cytotoxic compounds currently used in chemotherapy also bring about serious negative side effects in the patients (Firer and Gellerman, 2012). While monoclonal antibodies by themselves often have low therapeutic effects, they can be conjugated to highly potent drugs and act as their delivery vehicles to their target cells marked by specific surface antigens, preventing the drugs from adversely affecting healthy cells (Panowski et al., 2014). Especially promising are peptidic inhibitors that can target specific protein-protein interactions and bring about desired cellular changes, including apoptosis (Sorolla et al., 2020). Antibody drug conjugates (ADCs), as these pairings are known, have been lauded as a revolutionary step in anti-cancer drug development, but successful clinical application has proven challenging and the number of clinically approved systems remains low due to the complicated requirement of optimizing (Peters and Brown, 2015; Tang et al., 2019). Successful development of an ADC is a complicated process that requires the simultaneous optimization of tumour-specific antigen and its counterpart antibody, the stable linker between the antibody and the therapeutic agent, and finally, the agent itself. Current ADC development strategies often employ a limited number of existing microtubule inhibitors and their derivatives as the therapeutic agent payload, whose performances are measured with both *in vitro* and *in vivo* cytotoxicity tests (Lambert and Chari, 2014; Mullard, 2013). Efforts to expand the list of therapeutic agents that can be used as ADC payload to other classes of compounds are ongoing, but the choices still remain limited (Stein et al., 2018).

Given the requirements, novel ADC development would benefit from the availability of *in vitro* testing platforms through which the specificity and the efficacy of both the antibody and the drug can be evaluated. A number of human cell line models exists for testing cellular response to anticancer drugs, but ADC development entails some additional requirements to what is already available (Niu and Wang, 2015). Core requirements of a good testing platform for ADCs are that both the test and control cell types express the common antigens targeted by the antibodies at similar levels, and that any differences in the cellular changes induced by the drug stem mainly from the disruption of its target. If the first condition is not met, drugs would not be absorbed into the test and control cells at the same levels, complicating downstream efficacy evaluations. As for the second condition, it is necessary to limit the extent of off-target effects of the administered drugs so that they would not add layers of complication in interpreting the cellular changes introduced by the drugs. Unfortunately, because of the general lack of cell line pairings that differ in the drug target genes only, researchers have to contend with using two distinct cell types with differing origins to evaluate drug efficacy, despite the inherent risks of doing so. For example, there are numerous ongoing studies on p53, which is the most well-known and researched tumour suppressor gene, and a large portion of these studies focus on manipulating its expression in cancer cells through various therapeutic means including ADCs (Dangelmaier et al., 2019; Hafner et al., 2019; Kamp et al., 2016; Leroy et al., 2017; Mantovani et al., 2019). Another advantage afforded by well researched genes such as p53 is that one can utilize the already existing drugs and focus on the development of novel delivery mechanisms only. A popular testing platform for this purpose consists of the glioblastoma cell lines U87MG and U251MG, with the former having wild type p53 and the latter with an inactivated version that can no longer interact with MDM2, the main regulator of p53 expression (Cobbs et al., 2008; Leroy et al., 2017; Qin et al., 2019; Yount et al., 2001). The testing platform based on these two cell line pairs is particularly pertinent to ADC research, as they both express integrin ɑvβ3 that can be targeted by the same antibody and drugs that disrupt p53-MDM2 interaction are under intense research focus. Integrin ɑvβ3 is a frequently targeted cancer cell surface marker, as it is well known that its expression changes with tumour growth and metastasis (Sloan et al., 2006; Weigelt et al., 2005). However, although both U87MG and U251MG are glioblastoma cell lines, they are actually distinct cell types with completely different sources of origin, such that their genetic differences extend far beyond just p53 (Camphausen et al., 2005). Thus, even though many studies on p53 are performed on these two cell lines due to the known mutation in U251MG, the interpretation of the obtained results can be obfuscated by other inherent cellular differences.

Ideally, the best way to overcome this hurdle is to create targeted mutants of specific cell lines using genome engineering. One of the major developments in the field of genome engineering during the last decade has been the utilization of the CRISPR/Cas9 system for targeted gene editing, bringing unprecedented precision and ease of use (Hsu et al., 2014; Jinek et al., 2012; Ran et al., 2013). The system and its derivatives have gone through intense development and refinement, and they are now employed for a multitude of selective gene editing and regulation modifications. Wanzel *et al*. have employed this system to create p53 knockout mutants in the human colorectal cancer cell line HCT116 to investigate the mechanisms behind the disruption of p53-MDM2 interaction by the known inhibitor RITA (Wanzel et al., 2015). They targeted multiple intronic and exonic regions, resulting in mutants with both small indel mutations and large deletions and experimentally validated p53 inactivation in the resulting p53 knockout mutants. However, there was no comprehensive follow-up analysis of the p53 KOs to examine the extent of the transcriptomic changes induced by these mutations, both from the p53 modification or other potential off target effects.

In this study, we employed CRISPR-Cas9 to make targeted mutations of the *TP53* gene in U87MG, with the goal of creating a mutant cell line that has otherwise minimal differences from the wild type. We performed transcriptome analysis of the mutant clones created to examine the genetic differences using Cap Analysis of Gene Expression (CAGE), which can identify the location of transcription start sites and their expression levels (Carninci et al., 2006; Kanamori-Katayama et al., 2011). We confirmed that the extent of genetic differences between the wild type U87MG cells and the mutant clones were not extensive, and that these changes were mostly attributable to the inactivation of p53. We down-selected from the clones based on further analysis of their transcriptomic profiles compared to the wild type, and subjected them to further experimental validation to ascertain that there are no major changes to their *in vivo* phenotypes compared to the wild type U87MG cells and that other attributes necessary for their utility as testing platform for ADC development remain intact.

## 2 Materials and Methods

### 2.1 CRISPR-Cas9

#### Cell culture

U87MG cells (ATCC HTB-14) were grown in high glucose DMEM (Gibco) supplemented with 10% fetal bovine serum (Gibco), 100 U/mL penicillin (Gibco) and 100 μg/mL streptomycin (Gibco) at 37°C with 5% CO_2_.

#### Lentiviruses

To prepare the *TP53* knockout cells by CRISPR-Cas9 gene editing, sgRNA targeting TP53 gene exon 5 (5’-GTTGATTCCACACCCCC.GCCcgg-3’) was cloned into lentiGuide-Puro vector (Addgene, #52963), named lentiGuide-Puro_e5.2. Lentivirus for lentiCas9-Blast (Addgene, #52962) and lentiGuide-e5.2-Puro were prepared in a mixture of the following packaging constructs: pCMV-VSV-G-RSV-Rev and pCAG-HIVgp (provided by RIKEN BRC, Japan) according to the protocol (Suzuki et al., 2012).

#### Mutation

LentiCas9-Blast lentivirus transduction was performed with 8 μg/mL polybrane supplemented DMEM at 2 MOI for 24 h. Fresh DMEM without polybrane was used 24 h after transduction to avoid cell damage. After 48 hours, transduced cells were selected onto medium containing 15 μg/mL blasticidin, and incubated for 7 days. LentiGuide-Puro_e5.2 lentivirus was transduced into Cas9-expressing cells at 2 MOI, and selected by 1 μg/mL puromycin and incubated for 4 days. Cleavage efficiency was estimated through indel detection performed using the Guide-it Mutation Detection Kit from Clonetech.

#### Clone selection

Individual clones were scaled up in a 10 cm dish, and screened with western blot using anti-human p53 IgG (CST, rabbit polyclonal). Sanger sequencing of the TP53 cleavage target region in the selected clones was performed using Applied Biosystems 3730xl DNA Analyzer with BigDye Terminator v3.1 Cycle Sequencing Kit (ThermoFisher Scientific), and aligned to the wild type region using CLUSTAL version 2.1 (Larkin et al., 2007).

#### Western blotting

Cells were lysed in RIPA buffer (Pierce) containing proteinase inhibitors (Halt Proteinase Inhibitor Cocktail, Thermo Scientific). Protein concentration was determined with BCA Protein Assay Kit (Pierce). Protein lysates (20 μg) were separated on NuPAGE Tris-Acetate Gels (Invitrogen) and transferred onto PVDF membranes. Membranes were blocked with 5% skim milk in TBST for 1 h at room temperature and incubated with anti-human p53 antibody (Cell Signaling Technologies, #9282, rabbit polyclonal, 1:1000) in TBST containing 5% skim milk (4°C, overnight). Membranes were incubated with anti-rabbit IgG-HRP (GE Healthcare, NA934V, 1:2000) in TBST for 2 h at room temperature. Detection was performed with ECL Prime (GE Healthcare) and imaged by chemiluminescence imaging system FUSION SOLO S (VILBER LOURMAT).

#### Immunofluorescence

Cells were seeded with 80,000 cells per well on a coverslip coated with L-polylysine into 24-well plates. After overnight incubation, cells were fixed with 4% paraformaldehyde in PBS for 10 min at room temperature. Cells were blocked with 1% BSA-0.3 M glycine in PBST and incubated with mouse anti-human integrin αvβ3 IgG (1:200, monoclonal, LM609, Abcam, ab190147) in 1% BSA-PBST overnight at 4°C. For isotype control, mouse (G3A1) IgG1 Isotype Control (Cell Signaling Technologies, #5451) was used. Cells were incubated with donkey anti-mouse IgG-Alexa 488 conjugate (1:200, Jackson ImmunoResearch Laboratories, code: 715-545-150) in 1% BSA-PBST for 1 h at room temperature. The samples were embedded in VECTASHEELD mounting medium (Vector Laboratories), and imaged by OLYMPUS IX71.

#### Reverse transcriptase polymerase chain reaction

With starting RNA amount of 100 ng per sample, reverse transcription of *ITGAV*, *ITGB3*, and *GAPDH* (internal control) was performed using SuperScript III and oligo-dT primers on GeneAmp PCR System 9700. Quantitative PCR was performed with Hotstar Taq, customized primers on the ABI 7900HT Fast Real-Time PCR System. Experiments were done in triplicates per cell type.

### 2.2 CAGE data generation and analysis

#### Data generation

Total RNA was extracted from samples using RNeasy Mini Kit (Qiagen). The CAGE libraries were constructed using the published nAnT-iCAGE protocol (Murata et al., 2014): 5 μg of total RNA per sample from the mutant clones and the wild type cells was reverse transcribed with SuperScript III reverse transcriptase to produce cDNAs, which were then biotinylated and cap trapped to capture their 5’ ends. After barcoding and purification, they were sequenced on illumina HiSeq 2500 (50 nt single read). The sequenced reads were processed using the MOIRAI pipeline (Hasegawa et al., 2014). After filtering for rDNA and low quality reads, they were mapped to the human genome (hg38) using BWA version bwa version 0.5.9 (r16), with default parameters except: (1) for bwa_aln, maximum long deletion (-d) = 10, best hit limit (-R) = 30, seed length (−l) = 32, and (2) for bwa_samse, maximum number of alignments (-n) = 100 (Li and Durbin, 2009). The reads were overlapped with the FANTOM5 robust promoter set (http://fantom.gsc.riken.jp/5/datafiles/latest/extra/CAGE_peaks/) and mapped to the nearest GENCODE v27 annotations within 500 bp (FANTOM Consortium, 2014; Frankish et al., 2019). CAGE clusters within 20 bp of each other were merged together, and enhancer clusters overlapping existing GENCODE annotations were removed. For each processed CAGE cluster, the number of overlapping 5’ ends of the sequencing reads were summed to arrive at the raw expression counts.

#### Data analysis

The expression counts table was first filtered for lowly expressed CAGE clusters by first converting the values to counts per million (CPM), then removing those expressed in less than 3 samples with at least 1 CPM and greater than 1 sample with at least 3 CPM. Differential expression analysis of the expression table was performed with edgeR (version 3.28.0) in R 3.6.1 with TMM normalization and TREAT function with log fold change threshold of 1 and false discovery rate threshold of 0.05 (R Development Core Team, 2008; Robinson et al., 2010). KEGG pathway enrichment was performed using edgeR’s built-in kegga function and filtered for p-value threshold of 0.05. Motif Activity Response Analysis (MARA) was performed with the implementation used in Alam *et al*., 2020 (Alam et al., 2020; Balwierz et al., 2014). Briefly, transcription factor binding sites were predicted using the SwissRegulon set of motifs in 300 bp downstream and 100 bp upstream regions around each CAGE cluster, and motif activities calculated with the normalized CAGE expression table (Pachkov et al., 2007). The motif activities were converted to Z scores, and their differences between each mutant clone and the wild type were taken and visualized. Motifs with Z scores greater than 2 were marked as the significant set of motifs with notable regulatory effects on expression. TP53 ChipSeq peaks data sets were downloaded from Chip-Atlas (https://chip-atlas.org/) and overlapped with FANTOM5 promoter regions with +/−500 window. The TF activities were then calculated as mean log2 fold change of expression for genes with TP53 binding at the promoters. Off target regions for the sgRNAs used were predicted using CasOFFinder (Bae et al., 2014), and the GENCODE v27 annotations within 1 kb in either direction were retrieved and compared against the list of differentially expressed genes. The scripts used to perform these analyses are available at (https://github.com/tjkwon/CRISPR_p53).

### 2.3 Cell proliferation assay

U87MG (p53 wild-type) or the p53-mutant cells were seeded into 96-well plates at a density of 2,000 cells/well, and incubated for 24 h before treatment. p53-MDM2 interaction inhibitor, or nutlin-3a was added to cells at different concentrations, and the cells were incubated for 5 days at 37 °C. Then, 10 μL of a WST-8 reagent (Nacalai Tesque, inc.) was added to each well, and the cells were incubated for 2 h at 37 °C. The absorbance was measured using a iMark™ Microplate Reader (Bio-Rad Laboratories, Inc.) at 450 nm to count the living cells. IC_50_ values for nutlin-3a were calculated from nonlinear regression analysis (four parameters) using GraphPad Prism 7 (GraphPad Software Inc., La Jolla, CA, USA).

### 2.4 Cellular uptake and intracellular localization

FITC-labeled RGD peptide (Gly-Arg-Gly-Asp-Ser-Pro-OH), or FITC-RGD peptide was purchased from AnaSpec (Fremont, CA, USA). To investigate cellular uptake of FITC-RGD peptide, U87MG or the p53-mutant cells were seeded into 24-well plates at a density of 2.5 × 105 cells/well, and incubated for 24 h before treatment. The cells were incubated with 20 μM FITC-RGD peptide for 4 h at 37 °C. After the cells were washed twice with PBS and harvested, the fluorescent intensity of cells was determined by Cell Sorter SH800 (Sony Corp., Tokyo, Japan) using SH800 software (Sony Corp.). To investigate intracellular localization of FITC-RGD peptide, U87MG or the p53-mutant cells were seeded into 3.5 cm glass bottom dishes at a density of 9 × 10_5_ cells/dish, and incubated for 24 h before treatment. Cells were incubated with 20 μM FITC-RGD peptide for 4 h at 37 °C. The cells were washed twice with PBS, and incubated with 100 nM LysoTracker Red DND-99 (Invitrogen, Thermo Fisher Scientific, Inc., Waltham, MA, USA) for 30 min at 37 °C. After washing twice with PBS, the cells were fixed in 4% paraformaldehyde for 20 min. After washing twice with PBS again, the coverslips were mounted on glass slides with ProLong™ Diamond Antifade Mountant with DAPI (Invitrogen, Thermo Fisher Scientific, Inc.), and observed using a confocal microscope (LSM710, Carl Zeiss GmbH, Oberkochen, Germany).

### 2.5 Tumour growth of rates in subcutaneously xenografted mice

Male BALB/c nu/nu mice (5-week old) were purchased from Japan SLC, Inc. (Hamamatsu, Japan). Mice were subcutaneously inoculated with U87MG or the p53-mutant cells (3.0 × 106 cells per mouse) in 100 μL of Hank’s balanced salt solution (HBSS, Nacalai Tesque, Inc., Kyoto, Japan). The tumour size was measured with a slide caliper, and tumour volume was calculated using the following formula: tumour volume (mm_3_) = 0.5 × length (mm) × [wide (mm)]_2_. All animal experimental protocols were approved by the Ethics Committee on Animal Care and Use of the RIKEN Kobe Institute.

For the above 4 assays, statistical differences were analyzed using one-way analysis of variance (ANOVA) followed by the Dunnett tests for multiple comparisons. Kaplan–Meier curves were generated, and log-rank tests were performed. *P* < 0.05 was considered to be statically significant.

## 3 Results

### 3.1. Generation of p53-mutated U87MG cells and initial comparisons to wild type cells

We decided on the CRISPR-Cas9 system based on guide RNA-containing plasmid transfection and NHEJ pathway, as the goal of this study was to simply mutate p53 leading to disruption of known downstream interactions, and as such did not require the specificity offered by other methods. To increase our chance of success, we initially attempted the mutation of the p53 gene body in U87MG cells with different sets of single guide RNAs (sgRNAs), targeting introns 1, 2 and 9 for large deletions and exons 3 and 5 for indels. When screened with Guide-it mutation detection kit and western blot, the sgRNA targeting exon 5 was found to have achieved successful indel mutations in the target region (Figure 1, Supplementary Figure 1). These indels were confirmed with Sanger sequencing, and 7 unique p53 mutant clones were identified; their lengths suggest that they likely introduced nonsense mutations (Supplementary Table 1). After culturing and scaling up the mutant clones, their morphology was visually compared against the wild type U87MG cells, and in general, they maintained similar shapes and growth rates as their wild type counterparts (Supplementary Figure 2). The only exception was clone c4.2, which exhibited a higher growth rate than the wild type cells. When immunostained for integrin αvβ3, all of the mutant clones exhibited fluorescence levels at similar levels to the wild type, suggesting that the expression of integrin αvβ3 was not impacted significantly. All of the mutant clones exhibited similar levels of the integrin expression, with no discernible differences in their fluorescence. We also carried out reverse transcriptase polymerase chain reaction (RT-PCR) of *ITGAV* and *ITGB3*, the two component genes of integrin αvβ3 (Supplementary Figure 3). All the mutant clones expressed *ITGAV* and *ITGB3* at levels similar or greater than that of the wild type except for clone c4.2, which showed slightly lower *ITGB3* expression, and similar trends could be observed when the expression levels were compared against *GAPDH* for internal control. These observations demonstrate that the unwanted negative regulation of the integrin genes from the CRISPR-Cas9 process was minimal. As we envision utilizing the created mutant clones as an ADC testing platform with integrin αvβ3 as the antigen, it is vital that the integrin expression is maintained in both the mutant clones and the wild type. Together, these results confirmed that the CRISPR-Cas9 process successfully disrupted p53 protein level abundance with minimal effects on cell morphology and integrin levels.

**Figure 1.**
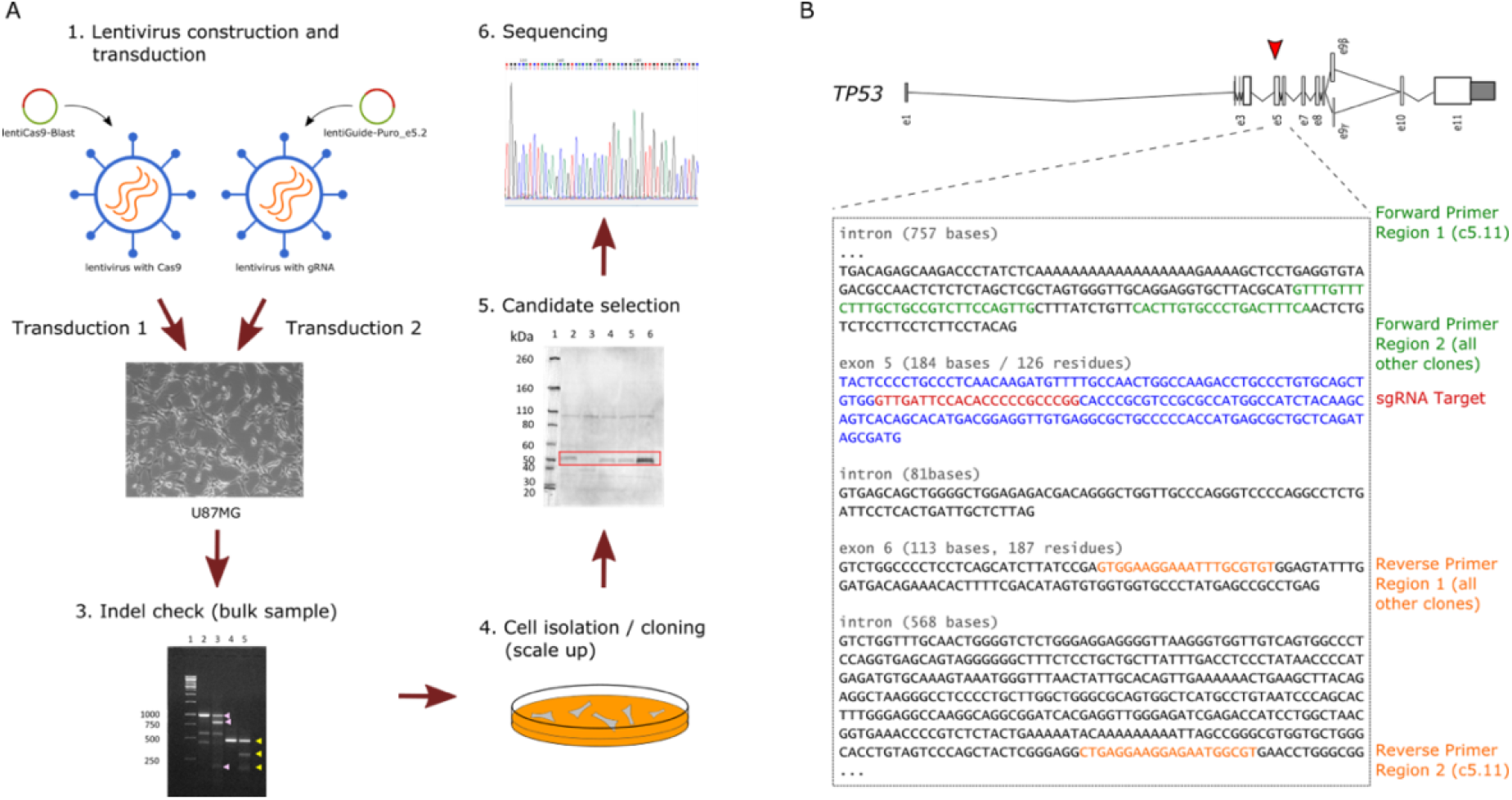
Targeted mutation of p53 gene at exon 5 in U87MG cells using CRISPR-Cas9 system. (a) Flowchart of the p53 mutation process. Virus particles containing the lentiCas9-Blast and lentiGuide-Puro_e5.2 vectors are constructed and transfected into U87MG cells in sequence. Those cells with indels are isolated and cloned for scale up, then candidates are selected based on western blot results, which are then subjected to Sanger sequencing for final confirmation of successful mutation. (b) The sgRNA target (red) in exon 5 (blue) and the forward and reverse primer regions for sub-cloning (green and orange, respectively) are shown. Clone 5.11 used different forward and reverse primer regions from the other clones.

### 3.2 Transcriptome comparison of mutants using CAGE analysis

p53 is involved in numerous cellular processes, and as such it regulates and is regulated by numerous interacting partners, both upstream and downstream. Thus, any mutation in p53 is bound to have both direct and indirect regulatory effects on many other genes. Furthermore, although the CRISPR-Cas9 system allows for highly specific targeted mutation, off-target effects resulting in unintended genomic changes still frequently occur and must be checked for. We wanted to evaluate the extent of these gene expression changes incurred by the p53 disruption using transcriptome profiling in each of the mutant clones and ascertain that the changes were acceptable for our purpose. We used cap analysis of gene expression (CAGE), which captures the 5’ ends of transcripts to identify transcription start sites (TSS) down to single nucleotide resolution, not only for promoters of protein coding genes but also noncoding RNAs and enhancers. Compared to RNA-seq, CAGE also has the advantage of detecting and analyzing alternative promoter usage at each gene. Comparing the transcriptome profiles of each mutant clone to that of the wild type using CAGE, enabled us to confirm the disruption of the p53-mediated pathways and use the measured expression changes to calculate their distances from the wild type cells.

We prepared the combined expression table for all samples by summing the number of reads per CAGE clusters based on the FANTOM5 promoter set (see Methods). After filtering for lowly expressed promoters and normalizing the expression levels, we arrived at the expression counts table of 19,931 CAGE clusters representing 14,994 genes (12,475 known) (Supplementary Table 2). With the normalized expression table, we first visualized the expression changes of *TP53*, *MDM2*, and *CDKN1A*, the core regulatory partners in the p53 pathway (Figure 2a) (Enge et al., 2009). While the expression levels of *TP53* did not show any consistent patterns across different clones, *MDM2* and *CDKN1A* showed consistent down-regulation in all of the mutant clones compared to the wild type. It is important to note that CAGE measures the transcription initiation events at the promoters only, and does not cover the entire transcript body as RNA-seq would. Since the sgRNAs targeted exon 5, the mutations did not result in abolishing of the transcription initiation events; in some clones, we even observed higher levels of expression than the wild type. Instead, the p53 transcripts produced in the mutant clones likely contain nonsense mutations that result in truncated proteins with reduced functionality. Conversely, we could clearly observe the effects of the mutation in the expression patterns of p53’s direct interacting partners MDM2 and CDKN1A: *MDM2* expression was reduced approximately by half, and *CDKN1A* showed greater than 3-fold decrease on average compared to the wild type. Other genes known to interact with p53 or MDM2 show down-regulation in the mutant clones as well (Supplementary Figure 4a) (Basu and Haldar, 1998; Bensaad et al., 2006; Meng et al., 2016; Serrano et al., 2013; Xiong et al., 2001). We also examined the expression levels of *ITGAV* and *IGTB3*, and confirmed that all mutant clones show expression levels similar to or greater than that of the wild type, as observed previously with RT-PCR. Especially, clone 4.8 showed notably elevated expression in *ITGAV* and *ITGB3*, suggesting that this clone may exhibit higher integrin presence on the cell surface than the other ones (Supplementary Figure 4b).

**Figure 2.**
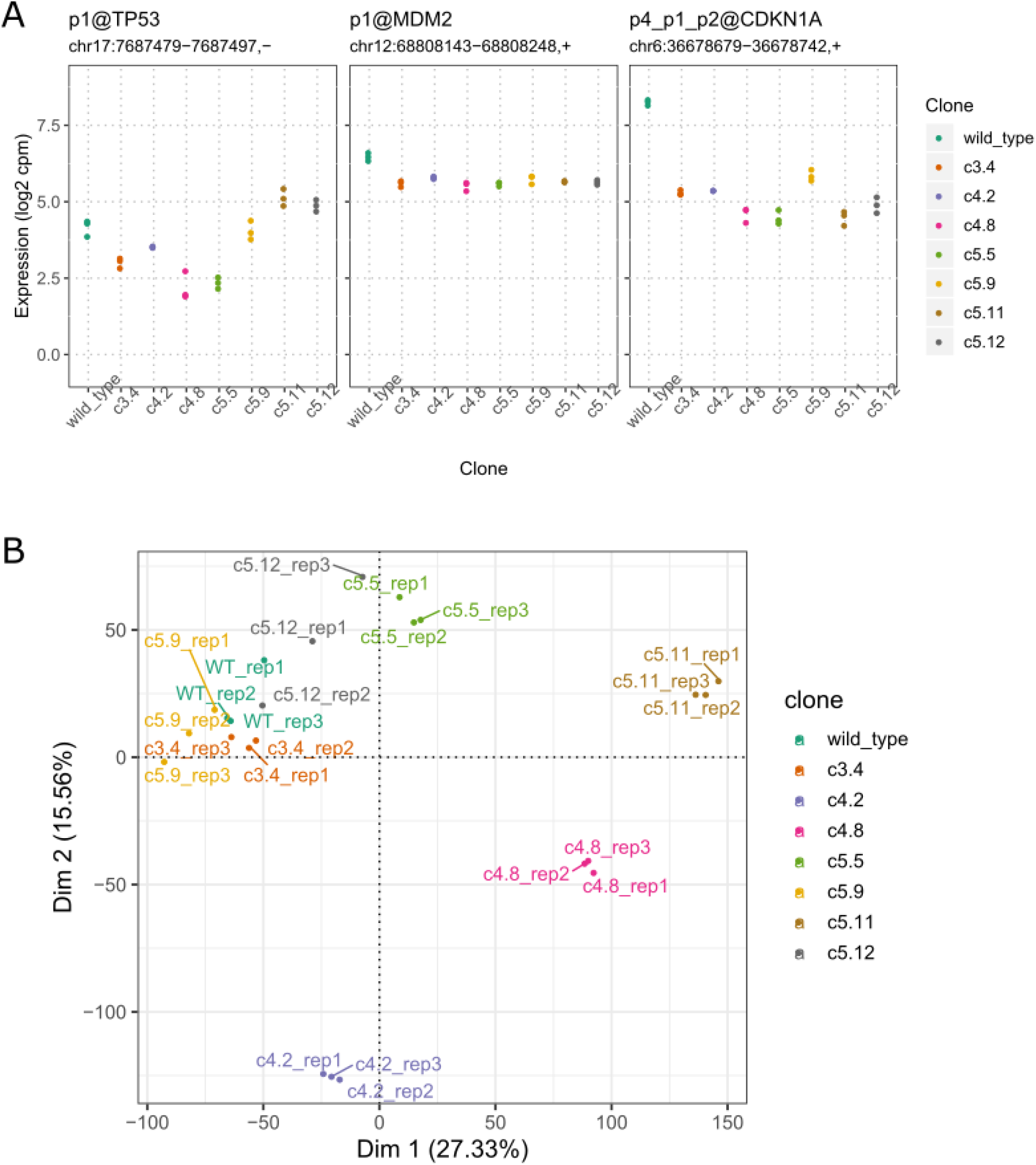
Comparison of the transcriptome profiles of the mutant clones against the wild type can be used to infer the degree of genomic changes caused by the respective mutation in each clone. (a) Normalized expression levels of *TP53*, *MDM2*, and *CDKN1A*. Both *MDM2* and *CDKN1A* exhibit down-regulation in the mutant clones. (b) PCA plot of the sample transcriptomes, with first two dimensions shown. Clones c3.4, c5.5, c5.9 and c5.12 cluster closely with the wild type.

With the confirmation that the key marker genes show consistent changes across the mutant clones, we proceeded to evaluate the overall transcriptome similarity of each mutant clone to the wild type using principal components analysis (PCA) (Figure 2b). By reducing the dimensionality of the entire transcriptome through orthogonal projection, PCA allowed us to visualize the proximity of different samples and infer their similarity to one another.

When visualized using the first two dimensions of the PCA, clones c3.4, c5.5, c5.9, and c5.12 clustered closely with the wild type, whereas the clones c4.2, c4.8, and c5.11 showed higher separation from one another as well as the other clones. As expected, the replicates of the same clone type formed the tightest clusters. Hierarchical clustering of the samples confirmed these results, with clones c4.2, c4.8 and c5.11 forming the outer branches of the clustering dendrogram (Supplementary Figure 5). The results suggest that the mutations in these ‘outer’ clones affected more widespread gene expression changes than the ‘inner’ clones, potentially leading to additional phenotype changes from the wild type such as the faster growth rate in clone 4.2 we observed.

To investigate the differences between clones in more detail, we performed differential expression analysis to identify the promoters in each mutant clone showing significant up or down regulation compared to the wild type expression levels (Supplementary Figure 6, Supplementary Table 3). The number of observed differentially expressed promoters were low in all clones. c5.5 showed the lowest number (37 up, 44 down) of both up- and down-regulated promoters, while c5.9 had the highest number of down-regulated promoters (417 down) and c5.12 the highest number of up-regulated promoters (126 up). The other clones showed similar numbers of differentially expressed promoters to one another, and overall there was no obvious relationship between the numbers of differentially expressed promoters and how they clustered together. This suggests that while numerous genes undergo minor but still significant expression changes due to the loss of p53 function, there may be a subset of genes with more drastic changes that contribute to the overall transcriptome differences to a higher degree. To examine whether the differentially expressed genes found were likely to be regulated by p53, we retrieved ChIP-seq binding data from ChIP-Atlas and analyzed whether there was an association between the differentially expressed genes and evidence of TP53 binding in their promoters (Supplementary Table 4). After determining which genes had TP53 binding evidence, we calculated the mean of the log fold change of expression of those genes in the mutant clones vs. the wild type and observed a consistent down-regulation of expression, further supporting the disruption of the p53-regulated genes resulting from its mutation (Figure 3a). We also made computational predictions of off-target regions affected by our sgRNAs and compared the results against our list of differentially expressed genes (Supplementary Table 5). Of the 32 candidate genes located within 1k base pairs from the affected regions, none were differentially expressed in the mutant clones vs. the wild type. The only overlap observed was from the list of differentially expressed genes between the fast growth clone c4.2 vs. the other mutant clones, where *TRABD2B* and *PDE7B* were found in our potential off-target gene list.

**Figure 3.**
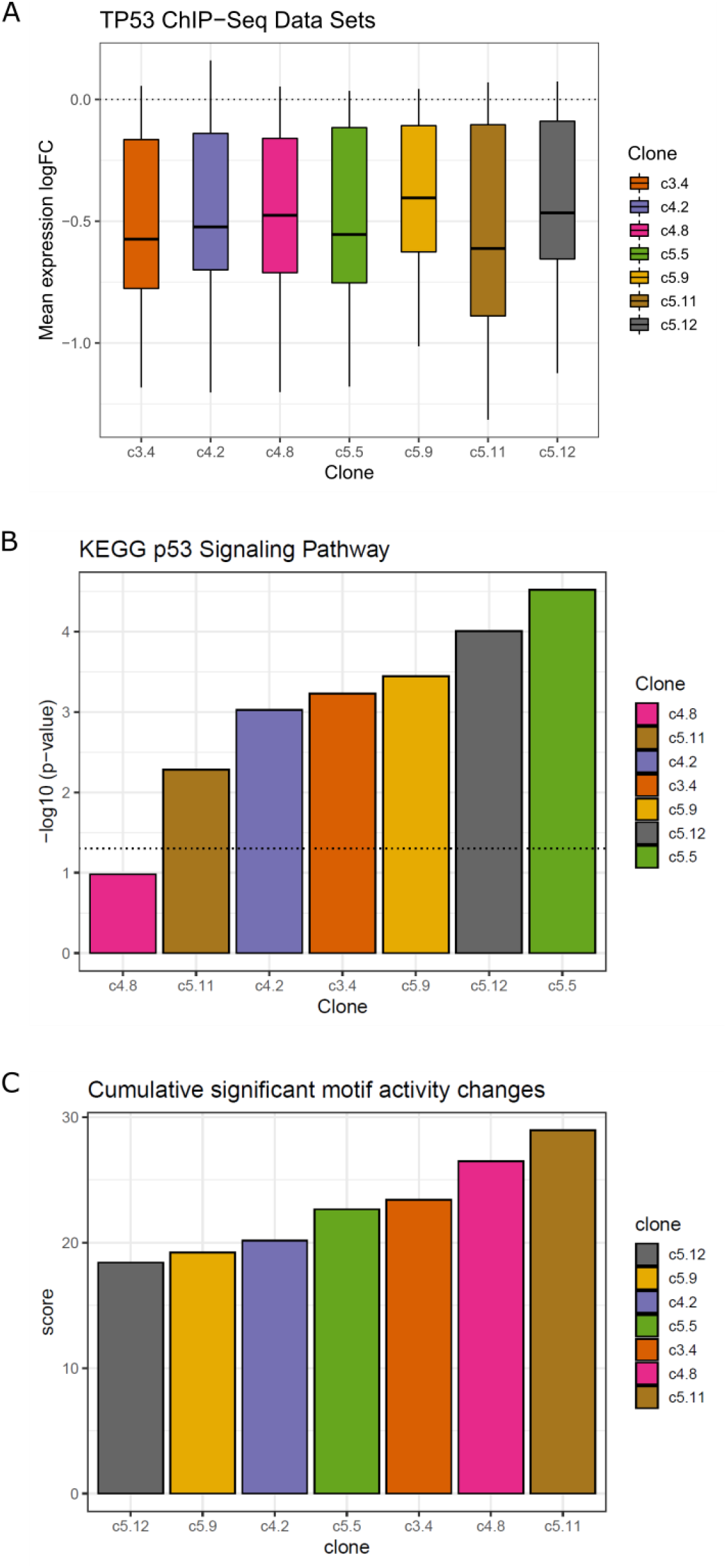
Down-regulated genes in the mutant clones vs. the wild type are enriched in p53 regulatory signals. (a) Genes with evidence of TP53 ChIP-seq binding show consistent down-regulation of expression across all samples compared to the wild type. The boxplot represents the mean log fold change of expression of the TP53-bound genes in mutant clones vs. the wild type. (b) Down-regulated genes show enrichment of the KEGG p53 signalling pathway. Barplot represents the negative log10 of p-values of pathway enrichment for each mutant clone. The clones are sorted in the order of increasing values, and the dotted line represents the p-value threshold of 0.05. (c) Greater motif activity differences between the ‘outer’ clones and the wild type suggests more extensive regulatory changes compared to the ‘inner’ clones. The differences in the motif activities between each mutant clone and the wild type for the significant set of motifs are summed and sorted to produce the overall ranking of the regulatory changes from the wild type.

In order to determine which cellular pathways are disrupted with these differentially expressed genes in each mutant clone, we performed functional analysis by identifying the enriched KEGG pathways in the list of differentially expressed genes, and examined in which group of genes the p53 signalling pathway was enriched in (Supplementary Table 6). As expected, we found that the genes contributing to the p53 signalling pathway were significantly enriched in the list of down-regulated genes among all comparisons (p < 0.05), except for the clone 4.8, which was placed away from the wild type samples in the PCA plot (Figure 3b, Supplemental Figure 7). When ranking the clones by the p53 signalling pathway enrichment scores (negative logarithm of enrichment p-values), we observed that the clones c4.8 and c5.11 scored the lowest while the clones c5.5, c5.12 and c5.9 scored the highest, in agreement with our previous observation from the PCA and hierarchical clustering. Further examining the top 10 pathways ordered by their median enrichment scores across the samples, we observed that the ‘inner’ clones show higher enrichment for cytokine-cytokine interaction pathways than the ‘outer’ clones, which is interesting as the link between p53 and inflammation has been previously reported (Figure 3b) (Gudkov et al., 2011). c5.5, which showed the highest enrichment for the p53 signalling pathway, showed elevated scores for other pathways as well, even compared to other ‘inner’ clones.

Finally, we performed Motif Activity Response Analysis (MARA) to probe the underlying regulatory factors behind the gene expression changes brought on by the mutations (Balwierz et al., 2014). MARA combines the predicted transcription factor (TF) binding sites (represented by TF motifs) found in promoters with the gene expression changes to calculate the motif activities, which can be used to estimate the responsible TFs for their regulation. When we applied the analysis using the SwissRegulon collection of TF motifs on our promoter regions (300 bp downstream and 100 upstream from each CAGE cluster), we detected a drastic decrease of TP53 motif activities in all the mutant clones compared to the wild type (Supplementary Figure 8, Supplementary Table 7). Besides the TP53 motif, those with significant activity variation included the motifs for NFY series of TFs and NFKB1/REL/RELA, key components of inflammation regulation (Ji et al., 2019). Another interesting motif was for PITX1, whose activity was down in the ‘outer’ clones c4.8 and c5.11, and is known to activate p53 by binding to its promoter (Liu and Lobie, 2007). Ideally, the clones selected for the purpose of this study should exhibit minimal regulatory changes from the wild type, except for TP53. To rank the clones based on this criterium, we first calculated the differences in motif activities between each mutant clone and the wild type, then summed these values for all significant motifs in each mutant clone, except for TP53. This creates a scoring system where the mutant clones with less regulatory differences would be assigned smaller values, and vice versa (Figure 3c). Consistent with previous observations, the ‘outer’ clones c4.8 and c5.11 obtained the highest scores, while the ‘inner’ clones c5.12 and c5.9 scored the lowest. One exception was c4.2, which obtained the third lowest score despite being an ‘outer’ clone.

### 3.3 Phenotypic comparison of selected clones to wild type

The transcriptome analysis confirmed that the major transcriptional changes in the mutant clones were associated with disrupted p53 function. However, it is clear that clone-specific transcriptomic differences also exist, with a subset of clones showing higher divergence from the wild type. To fully characterize how these differences will be manifested, we set out to perform further phenotype comparison experiments on the mutant clones. The phenotype comparison experiments performed in this study were designed to confirm 3 aspects of the mutant clones that would be crucial for the mutants to be useful for ADC development: disruption of TP53-MDM2 interaction, maintenance of cellular uptake and localization through surface integrins, and maintenance of tumorigenicity *in vivo*. Those mutant clones that meet all 3 criteria would qualify as the final candidates for ADC testing platform.

Before embarking on the experiments, however, we attempted to predict which clones were likely to show smaller or bigger phenotype changes compared to the wild type based on the transcriptome analysis results. We first separated out the clones c4.8 and c5.11 from the rest, as they exhibited consistently divergent behaviours from the wild type or other mutant clones. Clone c4.2 was not proximal to the ‘inner’ group clones in PCA, and obtained lower enrichment score for the p53 signalling pathway. However, its regulatory signal divergence from the wild type was at similar levels to those of the ‘inner’ group. The best performing clones were c 5.5, c5.9, and c5.12, but while c5.5 showed the highest enrichment for the p53 signalling pathway, it also exhibited higher regulatory changes than c5.9 and c5.12.

Taking these observations together, we decided to subject the following subset of mutant clones to further phenotypic studies and compare their behaviours against the wild type: c3.4, c5.5, c5.9, c5.12, and c4.2. We predicted that the clones c5.9 and c5.12 would prove to be the best candidates with the least phenotypic differences from the wild type. As for c4.2, while it did not appear to be proximal to the wild type, we were nonetheless interested in observing its phenotypic behaviour compared to the other clones, due to its conflicting transcriptomic signals and the previously noted rapid *in vitro* growth rate compared to other samples.

First, we wanted to confirm that the mutation in each clone led to successful disruption of the p53 protein’s capacity to interact with MDM2 and regulate cell growth. Nutlin-3a acts as an inhibitor of p53 and MDM2 interaction by blocking the p53-binding pocket of MDM2, and is known to activate the p53 pathway in cancer cells, leading to cell cycle arrest and apoptosis (Villalonga-Planells et al., 2011). Thus, we expected the wild type cells to have higher sensitivity to the inhibitory effects of nutlin-3a than the mutant clones, leading to slower growth rate at lower doses of the drug. After incubating the cells for 5 days with increasing dosage of nutlin-3a ranging from 0 to 100 μM, we performed colorimetric assay with WST-8 to measure the resulting relative cell concentration by taking the ratio of the absorbance against that of the no-treatment control group (Figure 4a). As expected, wild type U87MG cells showed high sensitivity to nutlin-3a, with decrease in cell number observed even at low doses of 0.1 μM, whereas the mutant clones required greater than 1 μM before any decrease could be observed. Clones c4.2, c5.5, and c5.12 showed the greatest resistance to the drug, requiring greater than 10 μM of dosage before significant decrease in cell counts could be observed. The IC50 values for nutlin-3a, which are half maximal inhibitory concentration representing the potency of an inhibitory drug, were more than 20-fold higher in the mutant clones than in the wild type cells (Figure 4b).

**Figure 4.**
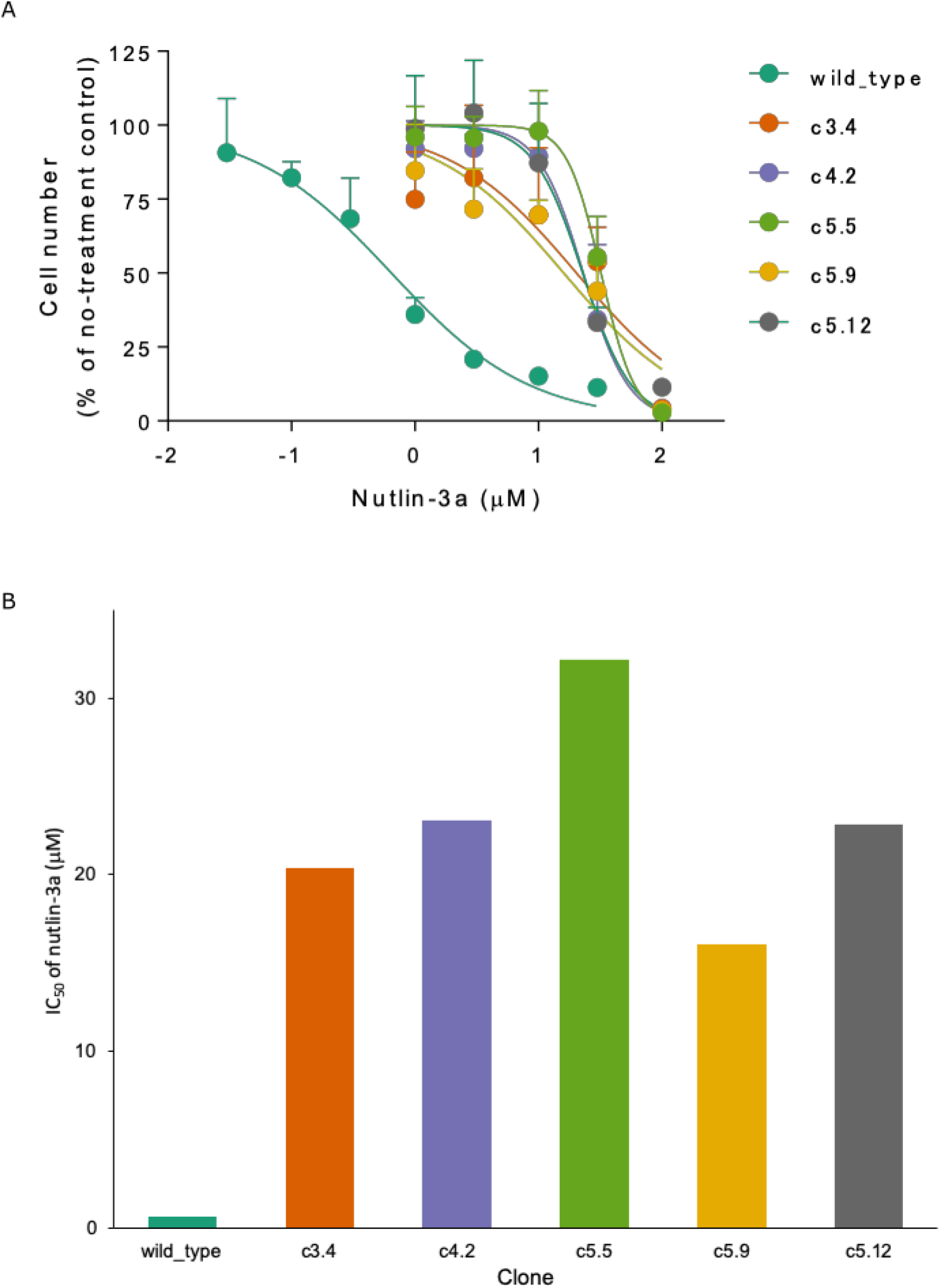
Cell proliferation assay results support successful disruption of p53-MDM2 interaction leading to decreased sensitivity to inhibitory drug nutlin-3a in mutant clones compared to the wild type. (a) Cell counts of nutlin-3a-treated groups with increasing drug dosage measured with WST-8 assay. Cells were incubated with nutlin-3a at 0 to 100 μM for 5 days at 37 °C. The count ratios of nutlin-3a-treated and non-treated cells in 4 replicates were plotted against different nutlin-3a concentrations (log scale). Results are expressed as the mean ± standard deviation. (b) IC_50_ values for nutlin-3a. All mutant clones show IC_50_ values > 20-fold over the wild type.

Next, we explored whether the mutant clones retained the same capacity for cellular uptake and intracellular localization of compounds that bind to the surface integrin ɑvβ3. As the goal of creating the mutants was to utilize them as testing platforms for ADC development, it was important to ascertain that not only was the integrin expressed at similar levels, as previously seen through fluorescence imaging (Supplementary Figure S2), but that their binding initiated the same level of endocytosis into the cells. As integrin ɑvβ3 is characterized by the exposed arginine-glycine-aspartic (RGD) tripeptide sequence, fluorescein isothiocyanate (FITC) can be conjugated to RGD peptides and induce its cellular uptake (Zheng et al., 2014). By performing cytometric analysis based on the fluorescence observed after the uptake, we could infer whether the activity of the expressed integrins is maintained across the mutant clones compared to the wild type U87MG. After incubation with FITC-RGD peptide, both the wild type cells and mutant clones showed fluorescence intensity shifts of greater than 90%, indicating they take up FITC-RGD peptides with similar efficiencies (Figure 5a). Clones c5.9 and c5.12 showed similar levels of mean fluorescence intensity (MFI) as the wild type (Figure 5b). Clones c3.4 and c5.5 showed small but still significant changes in the MFI levels from the wild type. c4.2, the outlier in our group, showed a significantly higher level compared to the rest of the samples. When we confirmed the localization of the fluorescence signals in the cells, they all showed punctuate patterns that co-localize with lysosome locations, suggesting that the FITC-RGD peptides were transported to lysosomes after their cellular uptake (Supplementary Figure 9).

**Figure 5.**
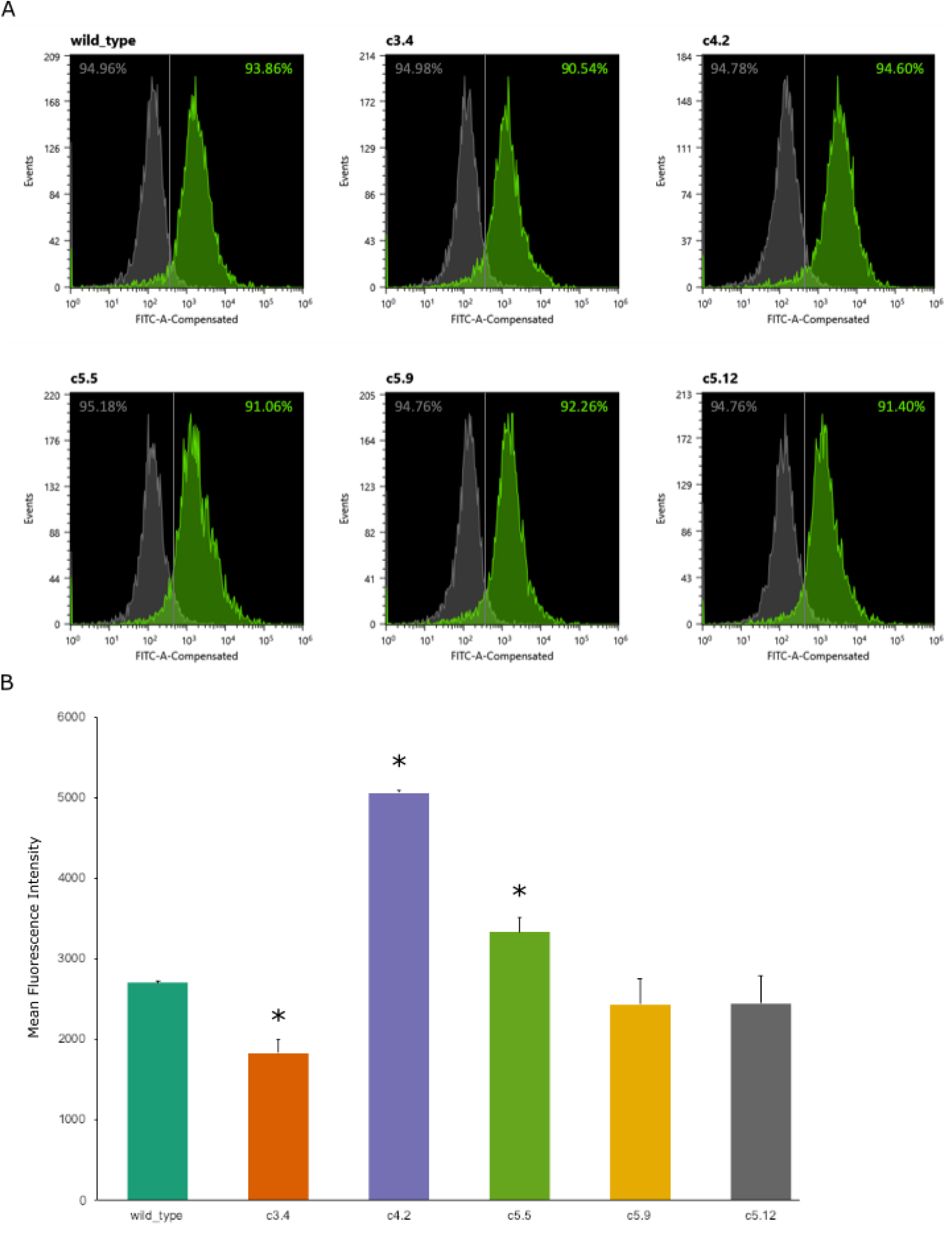
Flow cytometric analysis after incubation with FITC-RGD peptide shows similar levels of cellular uptake in both the wild type and mutant clones. The cells were incubated with the peptide at a final concentration of 20 μM for 4 h at 37°C. (a) Histograms of non-treated cells (gray) and FITC-RGD-treated cells (green). Vertical line represents the 95% mark of non-treated cells, and the fluorescein-positive ratios of cells can be seen on the right of the line. (b) Mean fluorescence intensity (MFI) of FITC-RGD-treated cells as an index of cellular uptake. The MFI values are calculated by subtracting the fluorescence intensities of the treated samples from those of the non-treated samples, and the results are expressed as the means ± standard deviations of 3 replicate samples. * *P* < 0.05 compared with the wild type group.

Finally, as we wanted to confirm the usability of the mutants for drug delivery system testing *in vivo*, we injected the cells subcutaneously into 5-week old mice and measured the tumour growth over the course of 70 days and examined whether there were any differences in their behaviour. Of the 5 mutant clones tested, clones c3.4, c4.2 and c5.12 achieved similar tumour growth as the wild type U87MG cells, while the clone 5.9 exhibited faster growth rate and c5.5 slower growth rate (Figure 6a). Most of the mutant clones and the wild type resulted in tumour growth in all the mice they were inoculated into, with the exception of clone c4.2 (7 out of 8 mice with tumour growth) and c5.5 (6 out of 8 mice with tumour growth) (Supplementary Table 8). One interesting observation was that the clone c4.2 showed similar *in vivo* growth as the wild type, even though *in vitro*, they exhibited higher growth rate. The same pattern of tumour growth rate differences could be observed when we measured the percentage of inoculated mice with tumour volumes greater than 1,000 mm3 over time (Figure 6b). Mice inoculated with clones c3.4, c4.2 and c5.12 exhibited similar rates of tumour growth, with greater than 50% of the mice exceeding 1,000 mm3 between 40 to 50 days after inoculation. Mice injected with the clones c5.9 and c5.5 differed significantly (*P* < 0.05), with those with clone c5.9 exceeding this figure before 30 days, while those with c5.5 needed more than 60 days.

**Figure 6.**
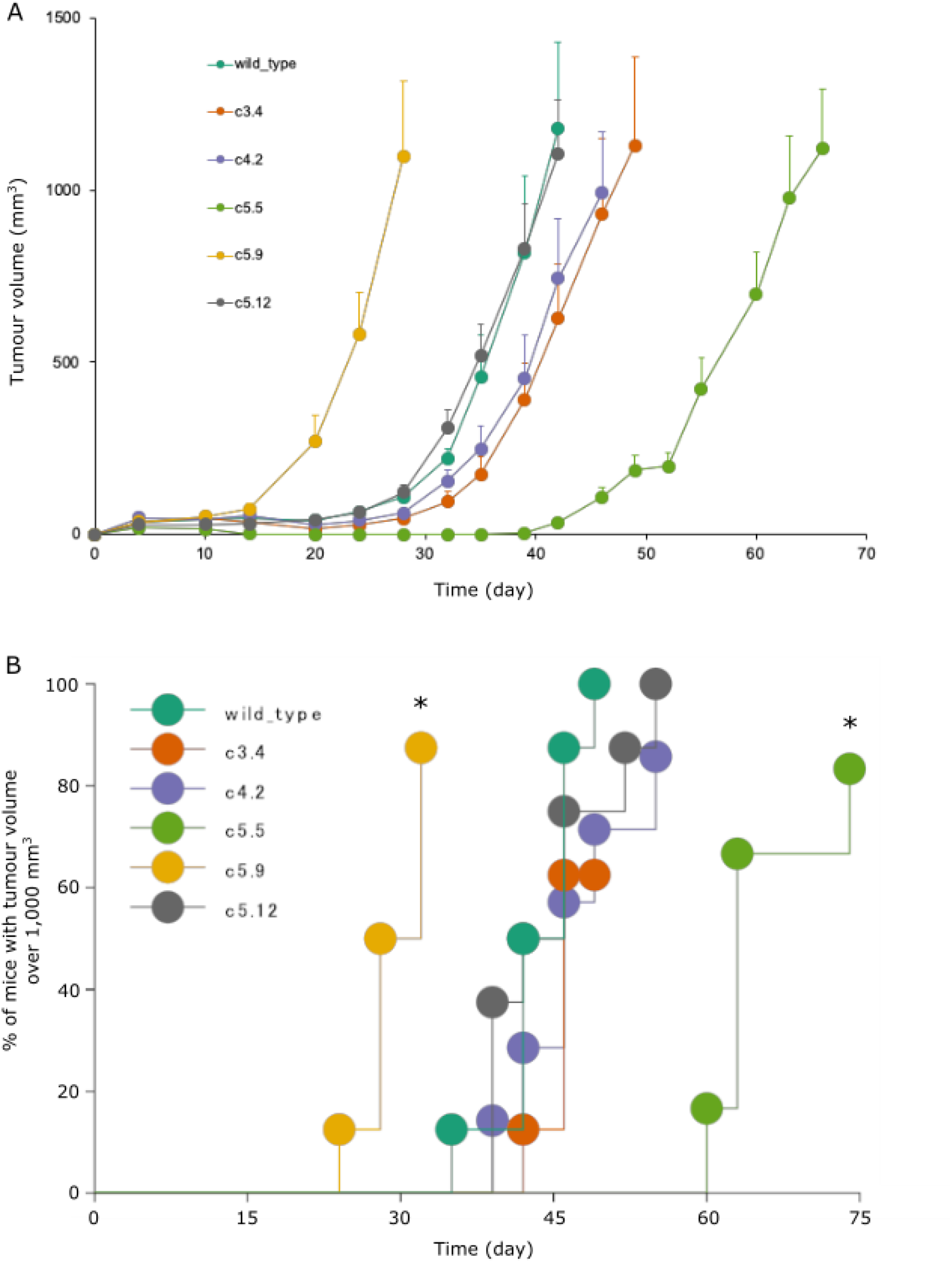
Tumour growth in BALB/c nu/nu mice after subcutaneous inoculation reveals the differences in the *in vivo* growth rates. (a) Increase in tumour volume over time. Results are expressed as a mean ± standard error of 8 mice (wild type, c5.9, c5.12 and c3.4), 7 mice (c4.2), or 6 mice (c5.5). Of the mutant clones tested, clones c3.4, c4.2 and c5.12 exhibit similar growth rates as the wild type. (b) Kaplan-Meier plots with the ratio of mice with tumour volume over 1,000 mm3 again show that the clones c5.5 and c5.9 exhibit different tumour growth patterns from the other samples. (* *P* < 0.05 compared with the U87MG tumour bearing mice group)

In summary, the phenotypic comparison experiments strongly support the successful creation of the p53 mutant clone c5.12 to be utilized for p53-specific *in vitro* testing platforms. This clone not only demonstrated disrupted interaction between p53 and MDM2 (Figure 4), but also maintained the capacity for cellular uptake through the surface integrin ɑvβ3 (Figure 5) and *in vivo* tumour growth as the wild type (Figure 6). While clones c3.4 and c4.2 also exhibited similar *in vivo* growth as the wild type, signs pointing to their significantly different cellular uptake prevented us from including them in the group of top candidate clones (Figure 5).

## 4 Discussion

In development of novel targeted therapeutics, there is an acute need for efficient procedures for establishing cell line-based testing platforms where the differences between the test and control samples are limited to the gene being targeted in order to avoid unwanted secondary effects stemming from other genomic differences. In order to improve upon the current situation where many researchers have to rely on two dissimilar cell lines, we employed CRISPR-Cas9 system to create targeted mutation of p53 in the U87MG cell line, while minimizing other genomic changes that may complicate interpretations of experimental results. While such endeavours have been previously made, we here took the extra step of performing detailed transcriptome profiling using CAGE and classified the clones into ‘inner’ and ‘outer’ groups, based on the amount of accompanying genetic disruptions compared to the wild type. Of the ‘inner’ group members, we further predicted that the clones c5.9 and c5.12 would be the best candidates to be used in the testing platform. We followed up with phenotype analysis experiments designed to fully characterize the mutant clones created and evaluate our predictions. The cell proliferation assay with p53-MDM2 interaction inhibitor nutlin-3a confirmed reduced interaction in the mutant clones. The FITC-RGD peptide uptake assay confirmed that the cellular uptake and localization through integrin ɑvβ3 remains intact in the mutant clones. Finally, tumour xenograft experiments in mice showed that 3 of the 5 mutant clones have similar *in vivo* growth rates as the wild type.

Based on the overall analysis results, we select the clone c5.12 as the best candidates for serving as the testing platform with the parent U87MG cell line as the control. Not only do this clone show relatively low genomic changes stemming from the p53 mutation, it is highly resistant to nutlin-3a, indicating strong inactivation of p53-MDM2 interaction, while still exhibiting similar levels of integrin functionality and *in vivo* growth behaviours. c5.12 was one of our two predicted candidates based on transcriptomic analysis. While c5.9, the other top predicted candidate, proved to have more rapid *in vivo* growth rate than the wild type, it performed comparatively similar in most of the other phenotypic measures.

This does not mean, however, that other clones are not of further research interests, as individual differences manifested through their transcriptome and phenotype changes present intriguing prospects of further investigation into their causes, and their inclusion in the development pipeline provides additional opportunities to control for clone specific effects. The CAGE technology used in this study offers many attractive advantages, as the data can be used to identify differentially expressed enhancers and various noncoding RNAs, both annotated and unannotated (Andersson et al., 2014; Hon et al., 2017). p53 is known to have complex regulatory relationships with many other genes, some of which are of noncoding type (Dangelmaier et al., 2019; Hu et al., 2018; Khan et al., 2019; Li et al., 2017). Some of these nonconding regulatory partners of p53 may be included in the list of differentially expressed promoters and enhancers we have identified in this study, which would serve as interesting research directions to follow upon.

Moving forward, we believe that we can employ the strategy used in this study for tailor-building of control-mutant cell line pairs targeting many different genes, both for the surface marker and the intracellular target. Being able to build a repertoire of these cell line platforms would greatly facilitate the ongoing development efforts for novel targeted therapeutics and their delivery methods. The newly created mutant clones are available as a resource, and can be provided upon request. The transcriptomic profiles of the clones in the form of CAGE data can be downloaded from GEO series **GSE155461** for those interested in further analysis.

## 5 Conclusion

Using the CRISPR-Cas9 system, we have successfully performed targeted mutation of TP53 in U87MG for the purpose of building an *in vitro* drug testing platform, and through transcriptomics and phenotypic studies, demonstrate that only limited biological changes from the wild type are found.

## Author Contributions

Conceptualization, EA, HM, ShT, HS; methodology, AK, KM, SaT, TA, MT, BK; software, AK, BK; investigation, AK, KM, SaT, TA, MT; validation, SaT, TA, KM, MT; formal analysis, AK, KM, BK; data curation, AK; writing—original draft preparation, AK, KM, SaT, TA; writing—review and editing, AK, KM, SaT, TA, BK, HS, ShT, HM, EA; visualization, AK, KM, SaT, TA; supervision, EA, HM; project administration, AK, EA, MF; funding acquisition, EA, ShT, HM, HS, MF.

## Acknowledgments

The authors would like to thank Piero Carninci, Mikako Shirouzu, Kensaku Sakamoto and Yasuyoshi Watanabe for creating the research environment that enabled this research.

