## Supplemental Information for "Efficient Development of Platform Cell Lines Using CRISPR-Cas9 and Transcriptomics Analysis"

### Supplementary Figures

**Figure S1**. Screening for successful mutant clones of CRISPR/Cas9 process targeting TP53 in U87MG cells. (a) Guide-it mutation detection kit confirmed indel mutation in exon 5 of TP53, with estimated cleavage efficiency of ~57%. (b) Based on western blot analysis of U87MG and e5.2 clones, 7 candidate mutant clones were selected for further analysis.

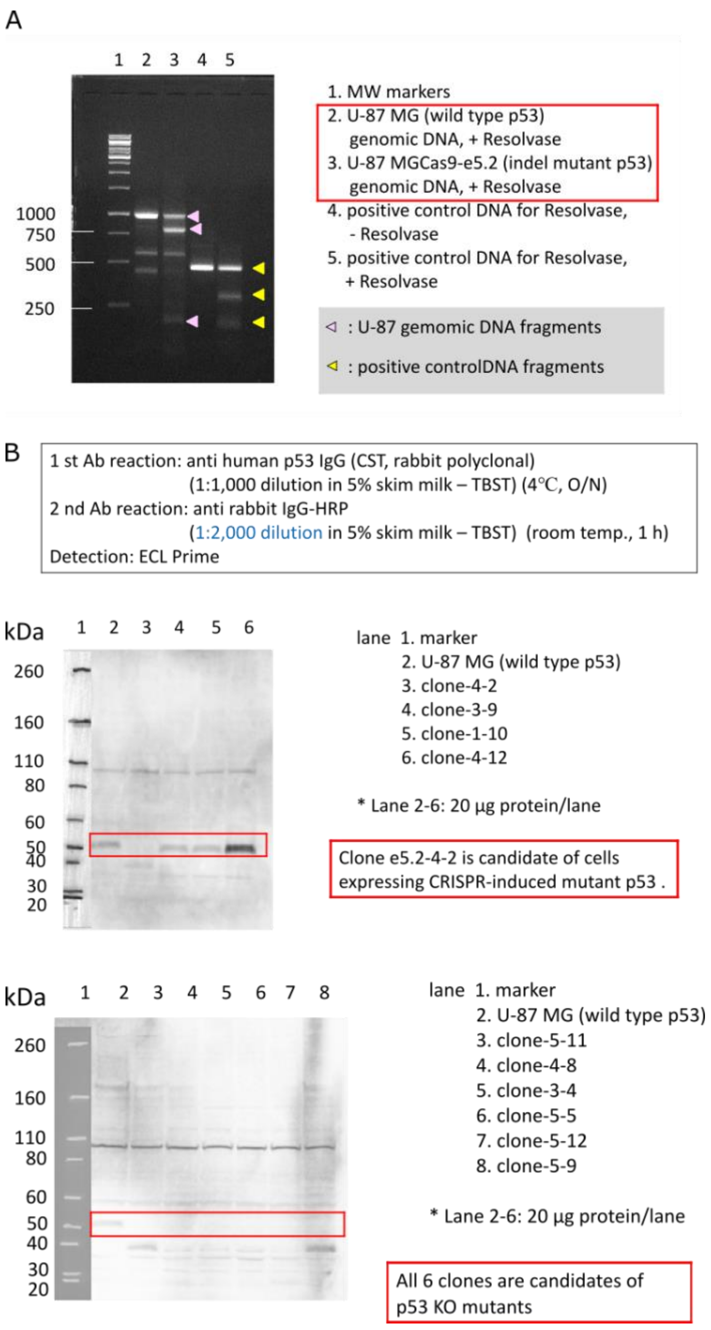

**Figure S2**. Cell morphology and integrin αvβ3 immunostaining of U87MG and p53 mutant clones. All mutant clones maintained similar morphology to the wild type and also exhibited similar levels of fluorescence for the integrin. (Top = transmission / Bottom = fluorescence)

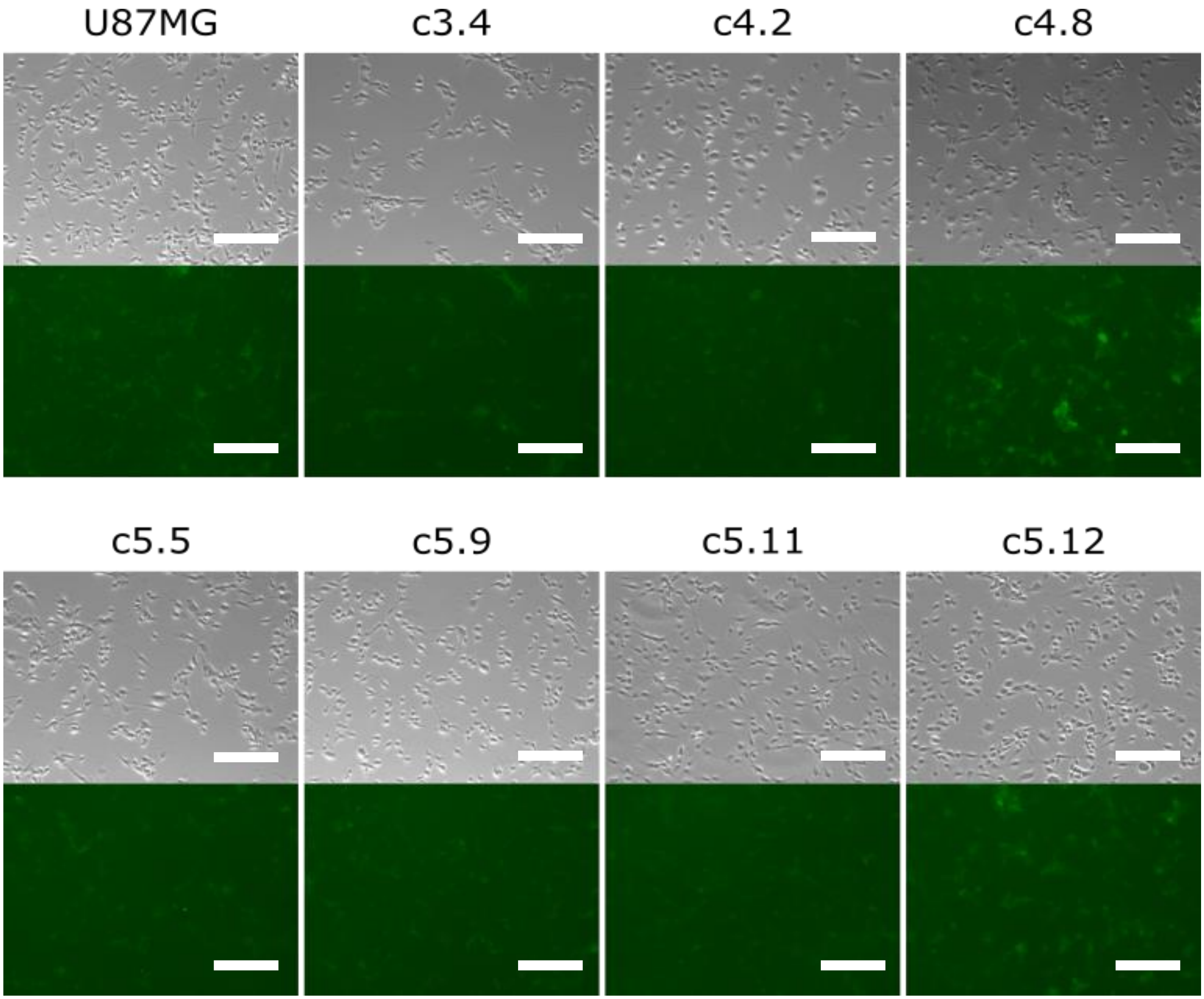


Bar = 200 μm

**Figure S3**. Expression levels of ITGAV, ITGB3 and GAPDH in p53 mutant clones and U87MG cells as measured by RT-PCR.

(a) ITGAV and ITGB3 expression levels after correction.

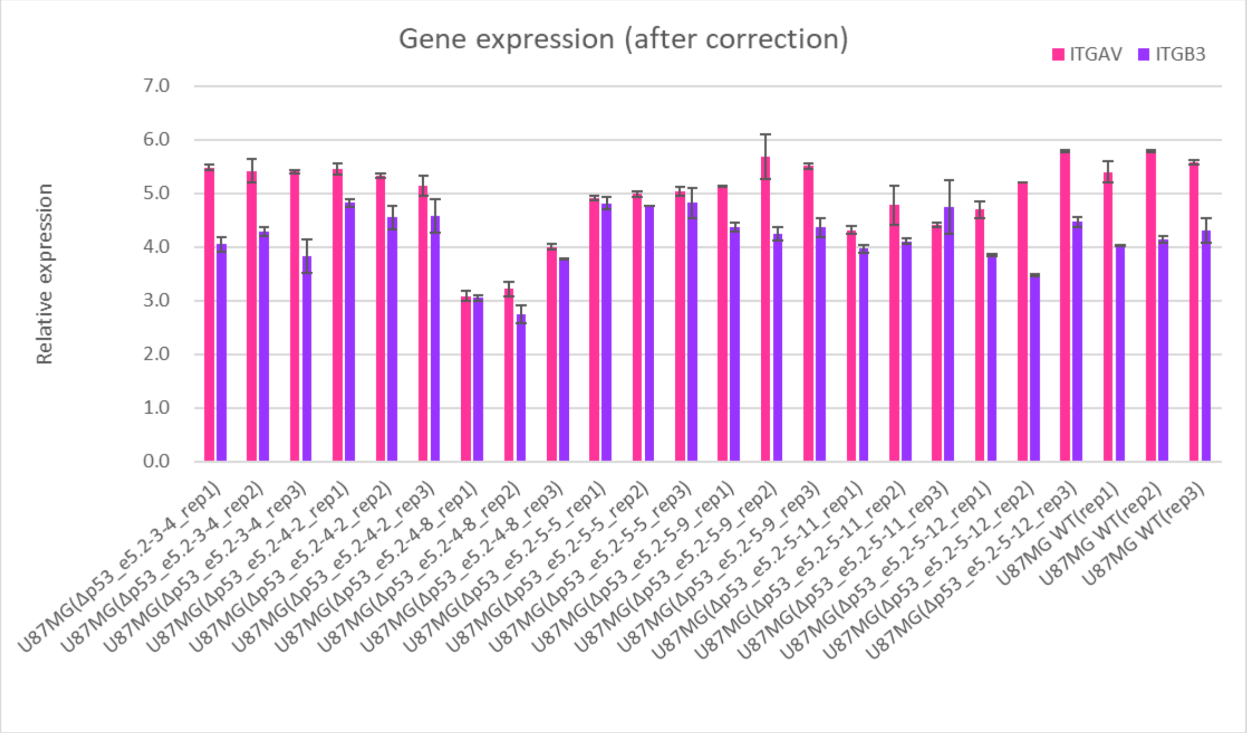


(b) Relative expression of ITGAV and ITGB3 with GAPDH as internal control.

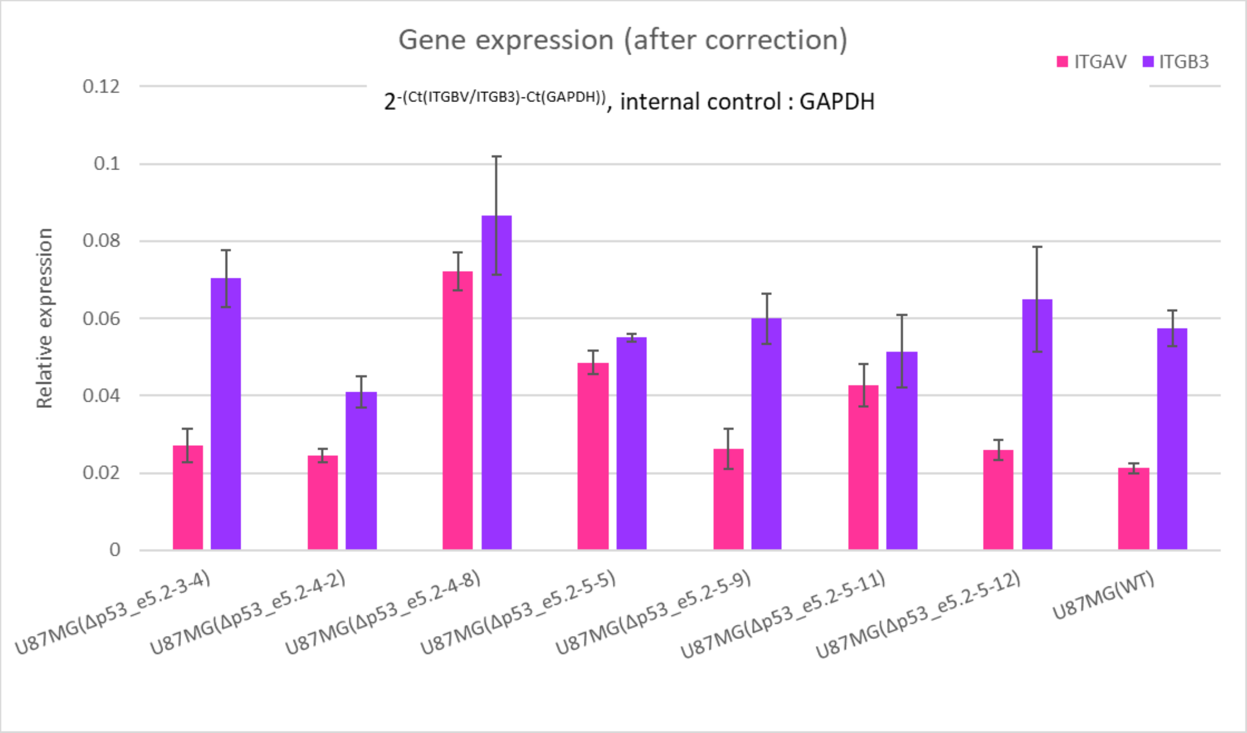

**Figure S4**. Promoter expression levels for marker genes from CAGE. Expression levels are given in counts per million.

(a) Genes known to interact with p53.

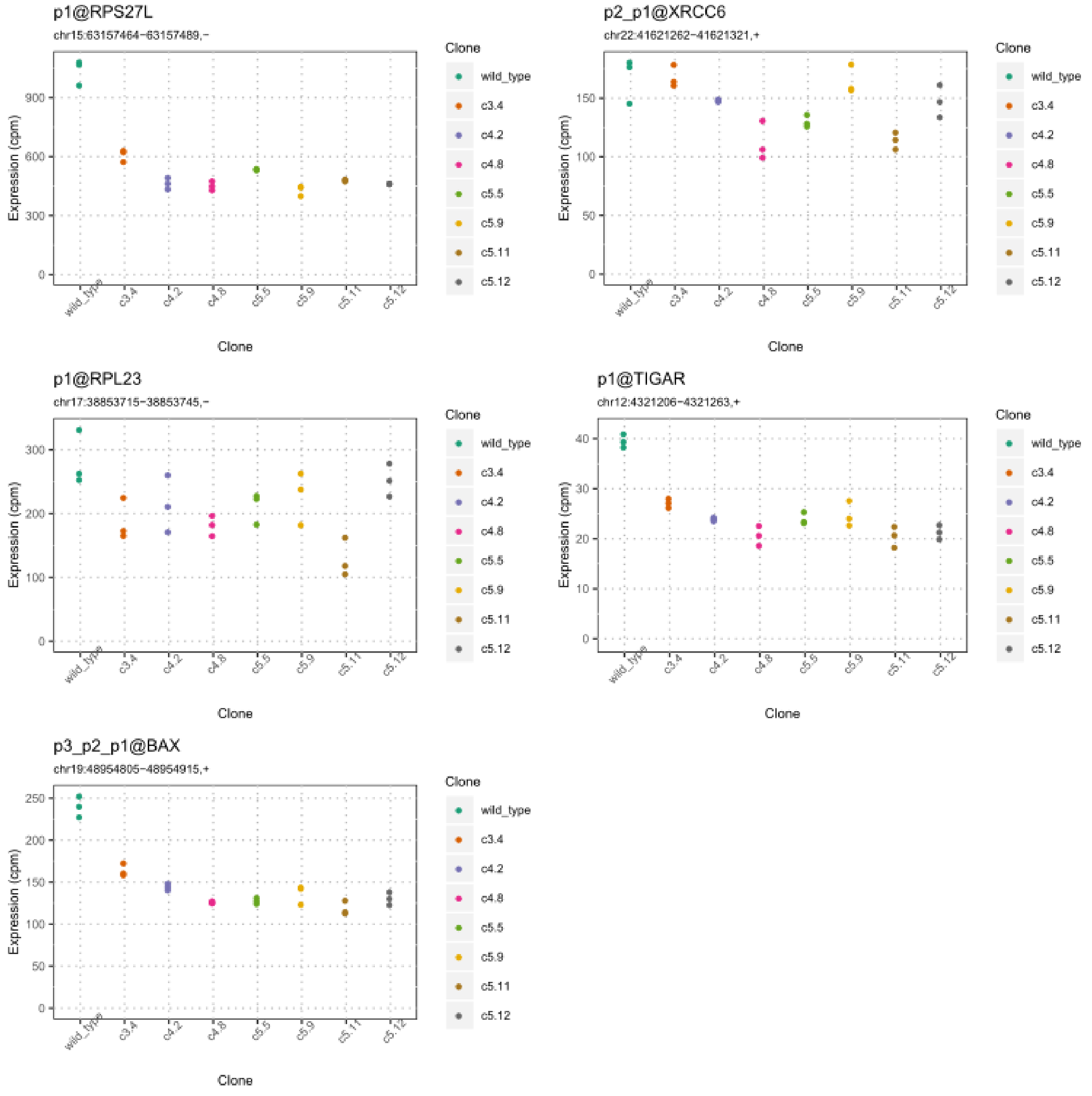


(b) ITGAV and ITGB3

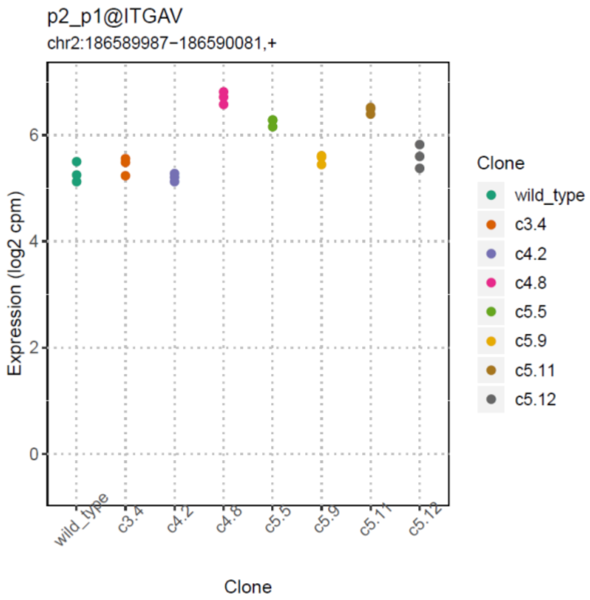
p2_p1@ITGAV
chr2:186589987−186590081,+
Expression (log2 cpm)
Clone
wild_type
c3.4
c4.2
c4.8
c5.5
c5.9
c5.11
c5.12

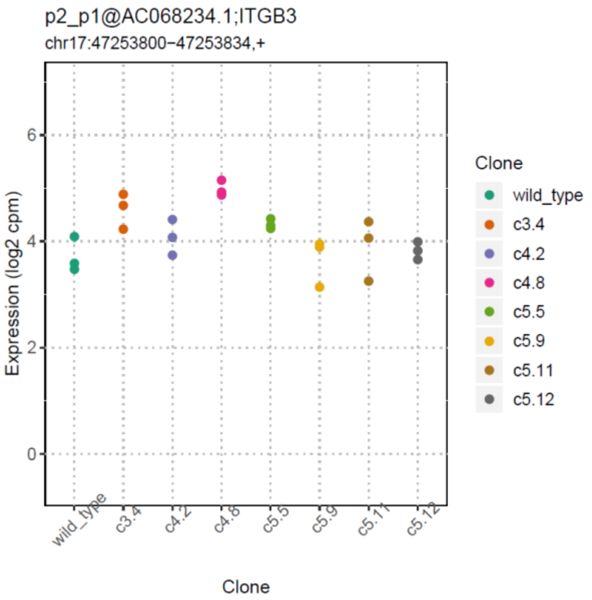
p2_p1@AC068234.1;ITGB3
chr17:47253800−47253834,+
Expression (log2 cpm)
Clone
wild_type
c3.4
c4.2
c4.8
c5.5
c5.9
c5.11
c5.12

**Figure S5**. Hierarchical clustering dendrogram of the samples based on CAGE expression data. Clones 5.11 and 4.8 are placed in the outermost branches of the tree.

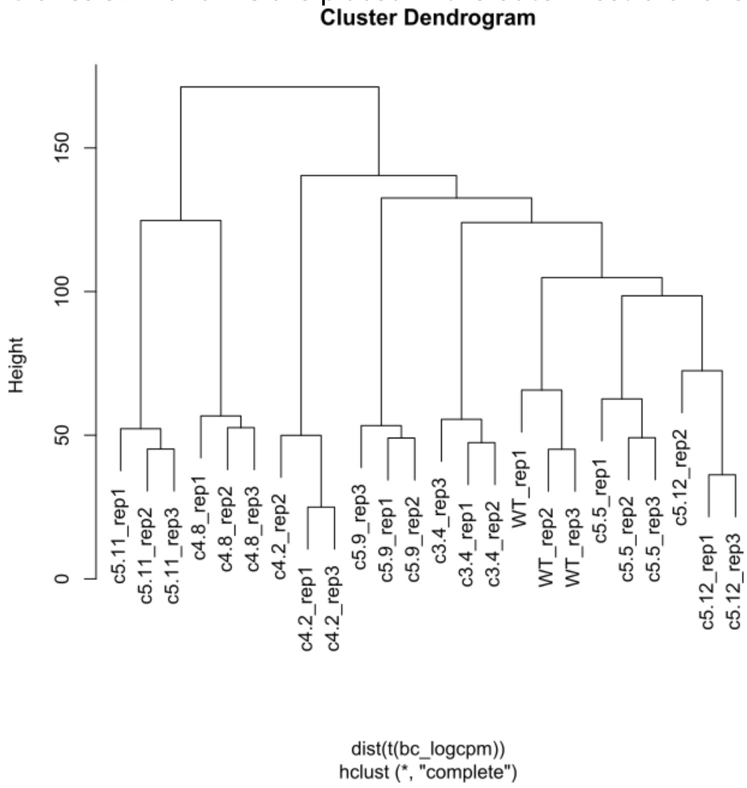

**Figure S6**. Smearplots of logFC vs. average logCPM of CAGE promoters between each mutant clone and the wild type samples. Red dots represent the promoters that are found to be significantly differentially expressed. ‘all.vs.wild_type’ represents the comparison of all mutant clones against the wild type, and ‘fast.vs.all’ represents the clone 4.2 vs. all other mutant clones (excluding wild type).

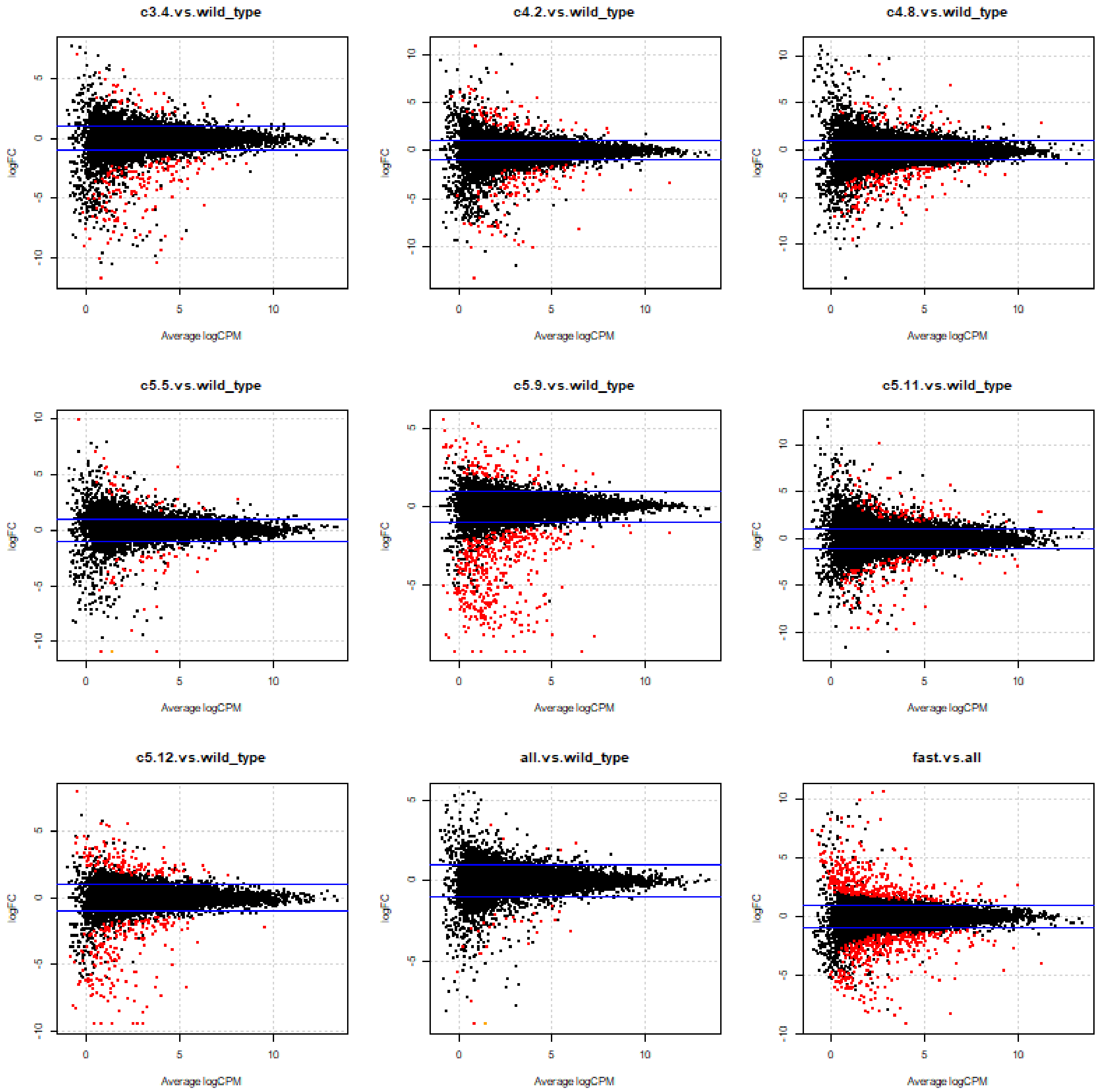

**Figure S7.** Heatmap of top 10 KEGG pathways enriched in genes down-regulated in each mutant clone vs. the wild type. The 'outer' group clones do not show as much enrichment in p53 signalling pathway and cytokine-related pathways as the 'inner' group clones do.

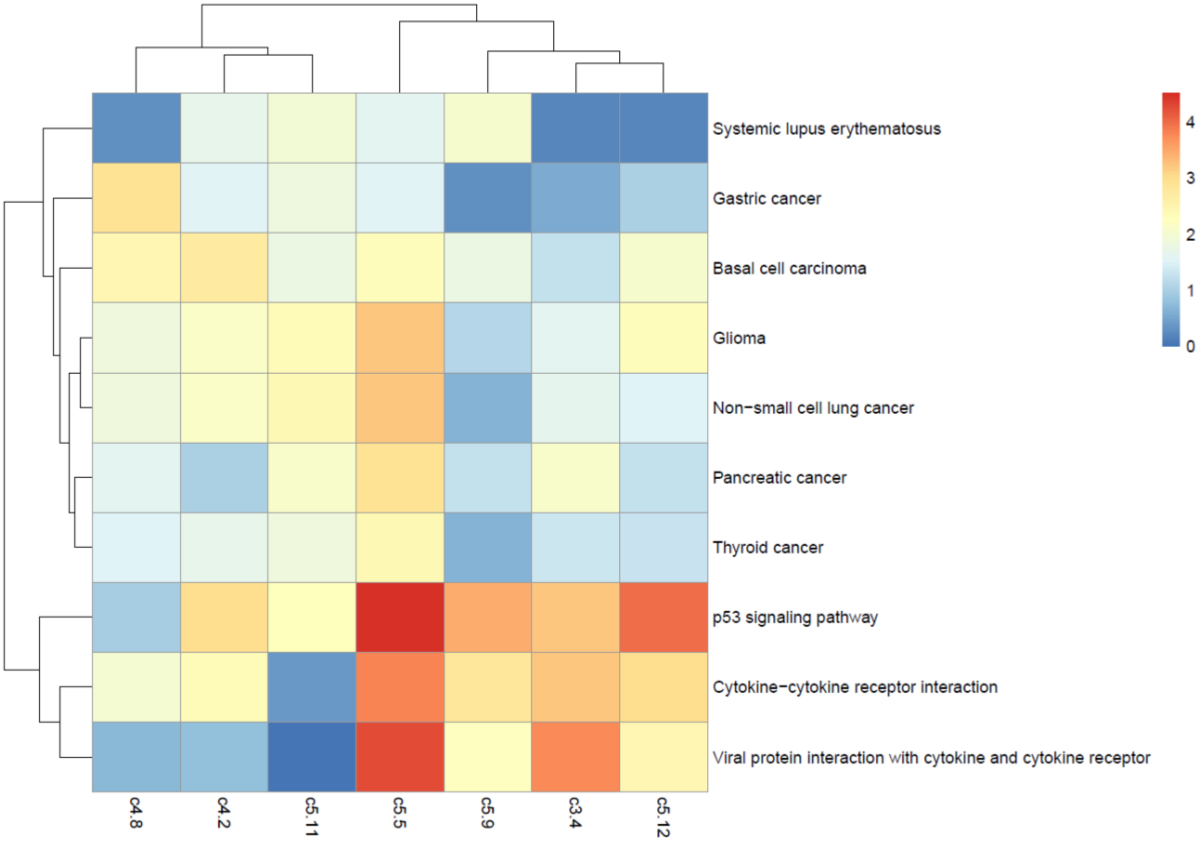

**Figure S8.** Heatmap of significant motif Z score differences between the mutant clones and the wild type. TP53 motif activity is down across all mutant clones compared to the wild type.

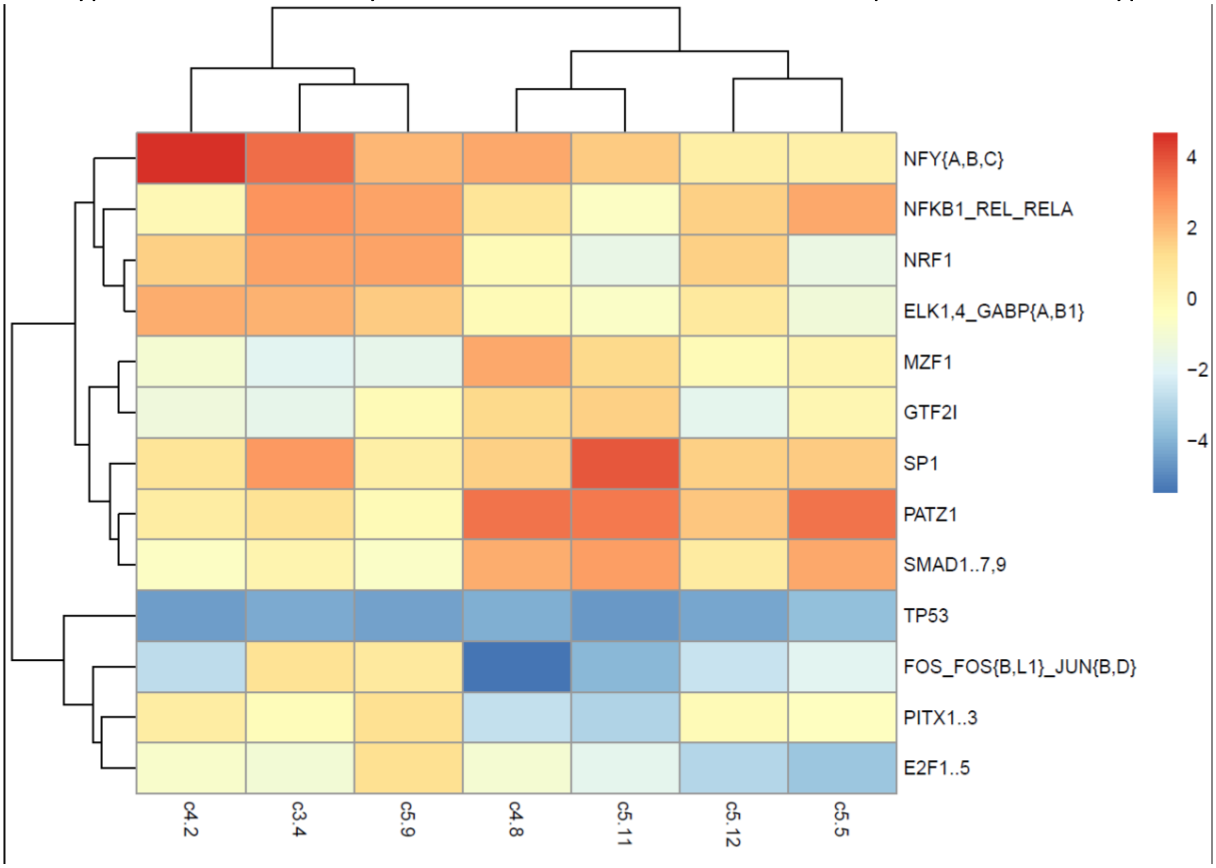

**Figure S9**. Localization of FITC-RGD peptide in U87MG and the p53 mutant cells. Fluorescent images after incubation with FITC-RGD peptide at final concentration of 20 M for 4 h at 37 C. Cell nuclei or lysosome was counterstained with DAPI or LysoTracker Red DND-99, respectively. All images were observed in a 1.0 m pinhole plane by confocal microscopy. White bars were 20 m. In colocalization images, yellow indicates colocalization of FITC-RGD peptide and cell lysosome.

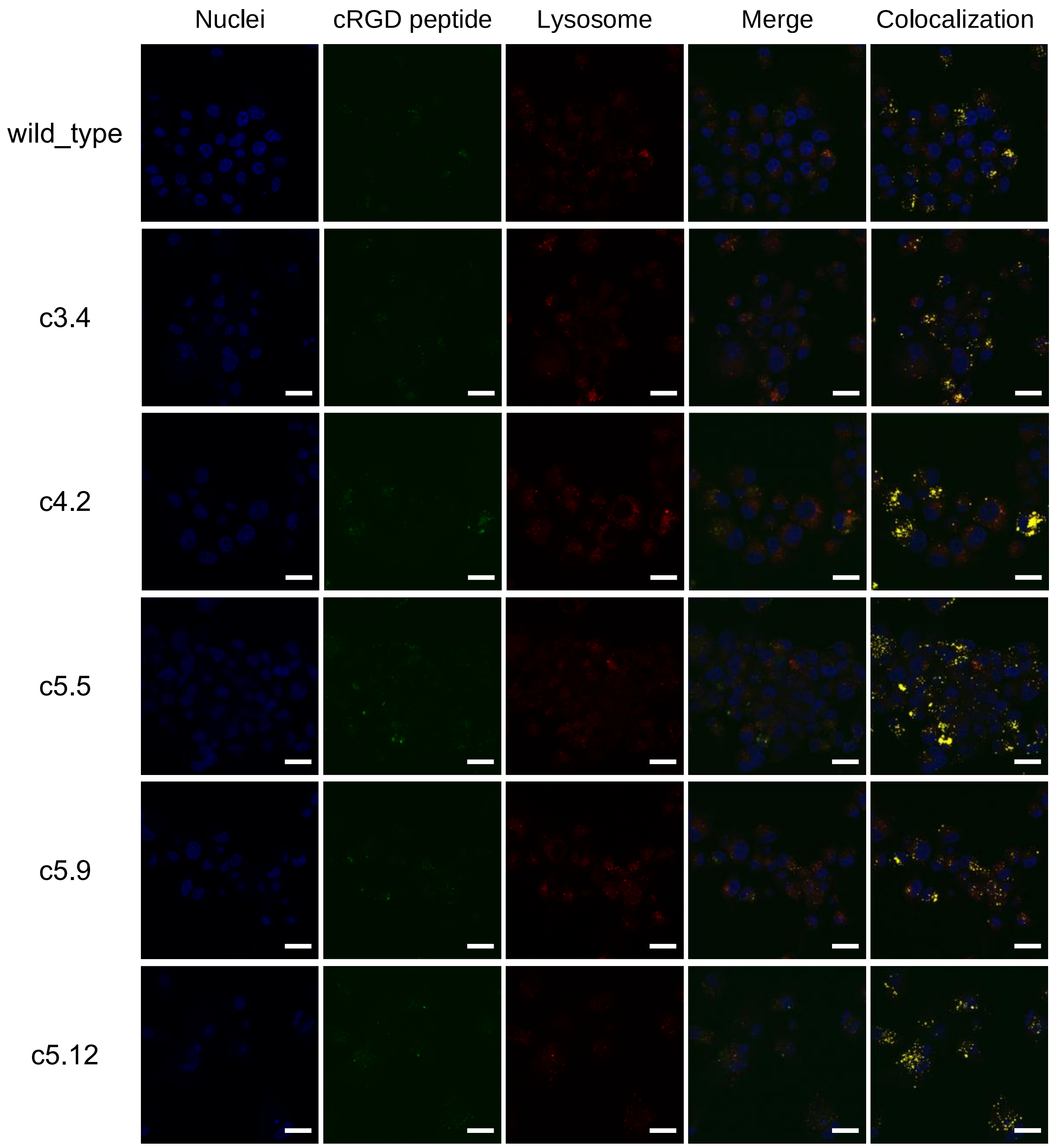

### Supplementary Tables

Table S1. Sanger sequencing results for mutant sequences. Tab 'total' summarises the results and identifies the specific mutation in each clone. (Table_S1.mutant_sequences.xlsx)

Table S2. Normalized and batch corrected CAGE expression data for all samples. Expression values given in log2 of counts per million. (Table_S2.batch_removed_log2_cpm.txt.gz)

Table S3. Number of differentially expressed CAGE clusters. (Full tables in Table_S3.DiffExp.zip)

a) All CAGE clusters

| clone | c3.4 | c4.2 | c4.8 | c5.5 | c5.9 | c5.11 | c5.12 |
|---|---|---|---|---|---|---|---|
| up | 36 | 68 | 51 | 37 | 93 | 81 | 126 |
| down | 157 | 106 | 140 | 44 | 417 | 110 | 188 |

b) Unique genes

| clone | c3.4 | c4.2 | c4.8 | c5.5 | c5.9 | c5.11 | c5.12 |
|---|---|---|---|---|---|---|---|
| up | 32 | 62 | 51 | 33 | 82 | 81 | 115 |
| down | 145 | 98 | 131 | 43 | 393 | 101 | 180 |

Table S4. Mean log fold change of expression in mutant clones vs. the wild type for genes with ChIP-seq binding evidence according to ChIP-Atlas. Each row represents the individual ChIP-seq data set. (Table_S4.chip_atlas_act.csv.zip)

Table S5. Off target region predictions by Cas-OFFinder for the sgRNAs used. The first 3 columns represent the predicted regions by Cas-OFFinder, and the columns start_1k and end_1k represent those coordinates extended by 1 kb. The 'gene' column represents the Gencode v27 gene annotation that overlapped with the extended region. (Table_S5.offtargets_1k_ext_genes.tsv)

Table S6. KEGG pathway enrichment analysis results for differentially expressed genes between the p53 mutant clones and the wild type U87MG. These are standard output files from edgeR's kegga function. (Table_S6.KEGG.zip)

Table S7. MARA results for the SwissRegulon set of motifs. The row names are the SwissRegulon motifs, and the columns contain the motif activities in each sample (columns 1-24), followed by their standard deviations (columns 25-48). The final column represents the overall z-score. (Table_S7.MARA.zip)

Table S8. Number of tumour bearing mice after inoculation with U87MG or the p53 mutant clones.

| U87MG | c3.4 | c4.2 | c5.5 | c5.9 | c5.12 |
|---|---|---|---|---|---|
| 8 | 8 | 7 | 6 | 8 | 8 |
